# Bioengineered ciprofloxacin loaded chitosan nanoparticles for the treatment of Bovine Mastitis

**DOI:** 10.1101/2020.05.09.085563

**Authors:** Preeti Yadav, Preksha Gaur, Awadh B. Yadav, Z. I. Huma, Neelesh Sharma

**Affiliations:** Centre of Biotechnology, University of Allahabad, Pryagraj-211002, India; Sher-e-Kashmir University of Agricultural Sciences & Technology, Jammu-181102, India

**Keywords:** Mastitis, Ciprofloxacin, Nanoparticles, drug delivery

## Abstract

Mastitis is the most devastating economic disease of dairy cattle. Mastitis in dairy cattle frequently occurs during the dry period or in early lactation. E. coli. and *S. aureus* is the main causative agent of mastitis*. S. aureus* has an ability to form encapsulated focuses in the udder and develop subclinical form of mastitis. This property of bacteria hinders the effective cure during lactation period. Antibiotics used for treatments have a short half-life span at the site of action because of frequent milking, therefore unable to maintain desired drug concentration for effective clearance of bacteria. We demonstrated the potential of ciprofloxacin encapsulated nanocarriers which can improve availability of drugs and could provide effective means of the treatment of mastitis. These drug loaded nanoparticles show low toxicity & slow clearance from the site of action. Antimicrobial studied against clinical strain of *E.coli.* and *Staphylococcus aureus* zone of inhibition depend on dose 0.5 mg to 2 mg/ml nanoparticles solution from 11.6 to 14.5 mm and 15 to 18mm. These nanoparticles show good antimicrobial activity in broth culture as well against bacteria.

## Introduction

Mastitis is economically the most important disease of dairy cattle. It reduced total milk production from 5-25% and in extreme cases 83%. The economic losses due to mastitis in US is 400-600 million dollar, while in India have augmented about 115 folds in the last five decades and presently it amounts to 995 million dollar per annum[1–3]. Mastitis in dairy cattle frequently occurs during the dry period or in early lactation. The majority of mastitis is of bacterial origin and just a few of species of bacteria account for most cases, such as *Escherichia coli, Staphylococcus aureus, Streptococcus uberis, Streptococcus dysgalactiae and Streptococcus agalactiae, Streptococcus bovis, Klebsiella pneumonia [4–6]. Staphylococcus aureus,* the main bacteria causing mastitis in dairy cows, has an ability to form encapsulated focuses in the upper part of the udder and develop a subclinical form of mastitis. This property of encapsulation hinders the bacteriological cure during lactation, because antibiotics used in this phase have short half-life in the target of action due to frequent milking and therefore unable to maintain therapeutic levels for long enough to determine the complete elimination of bacteria in cystic form. Nanocarriers have the possibility of providing controlled release of drug for a long time to maintain the desired level of drug at the infection site. Last few decades have seen unprecedented use of nanoparticles for drug delivery and various drug delivery platforms, especially liposomes, polymeric nanoparticles, dendrimers, and inorganic nanoparticles, have received significant attention. Drug interaction with nanocarriers through physical encapsulation, adsorption, or chemical conjugation have exhibited improvement in pharmacokinetics profile and therapeutic index. Persistent research efforts and encouraging results have resulted in a number of nanoparticles based drug delivery systems have been approved for clinical treatment of infectious diseases while many others are currently under various stages of pre-clinical and clinical tests[7,8]. Despite the presence of various preventive measures and management practices; there is an immediate need of effective therapeutics to treat mastitis. Chitosan is one of the most commonly used natural polymers in the production of nanomedicines because it displays very desirable characteristics for drug delivery and has proved very effective when formulated in a nanoparticulate form. Properties such as its cationic character and its solubility in aqueous medium have been reported as determinants of the success of this polysaccharide [9]. However, it’s most attractive property relies on the ability to adhere to mucosal surfaces, leading to a prolonged residence time at drug absorption sites and enabling higher drug permeation [9]. Chitosan has further demonstrated the capacity to enhance macromolecules epithelial permeation through the transient opening of tight epithelial junctions [9]. In addition, the polymer is known to be biocompatible and to exhibit very low toxicity [9], which are two mandatory requisites for drug delivery applications. Noticeably, chitosan has been referred to be more efficient at enhancing drug uptake when formulated in the nanoparticulate form, as compared to the solution [9].

Few studies from India, Pakistan, China and another part of the world have demonstrated high efficacy of ciprofloxacin against mastitis [9]. Ciprofloxacin (CPX) is a third generation of fluoroquinolone based antibiotics which show antimicrobials activity against positive and gram negative bacterium and their effective inhibition concentration is very small has been reported by different researcher group [10,11,5].

The nanoparticulate formulation administration by intramammary route is a promising alternative for the effective treatment of bovine mastitis. Therefore the present study was framed to develop intramammary nanoparticulate drug delivery system using chitosan (CS) as polymer because of its muco-adhesive and biodegradable properties and ciprofloxacin hydrochloride (CPX) as the model therapeutic drug because of its high efficacy against *S. aureus.*

## Materials and Methods

### Materials

*Ciprofloxin* gifted from, Chitosan, Sodium Triphosphate Pentabasic (TPP), Fluorescein Isothiocyanate (FITC) dye and Phorbol Myristate Acetate (PMA); was purchase from Sigma Bangalore India. Glacial Acetic Acid; NaOH; Dimethyl Sulphoxide (DMSO); Trehalose; Phosphate Buffer Saline (PBS); Syringe filter (0.22 μ pore size), RPMI culture media (Hi-media).

### Nanoparticles Preparation

The chitosan nanoparticles were prepared by ionic gelation method as previously described with slight modification [12]. Briefly, chitosan (1-2% W/V) was dissolved in 1% (V/V) glacial acetic acid solution and the pH adjusted to 5 using 1 N NaOH. Then, TPP solution (0.5-1 mg/ml; adjusted to pH 5 was added to the chitosan solution, and the mixture was stirred (750 rpm) for 30 min at room temperature. Drug-loaded nanoparticles were prepared by adding different amounts of Ciprofloxacin hydrochloride (CPX) (0.5, 1.0 and 1.5 mg/ml) into the chitosan solution and the above mentioned method was used. FITC dye loaded chitosan nanoparticles were prepared by adding 200 μl of FITC-DMSO (1 mg/ml) solution into the chitosan solution and polymerized with TPP in the dark to avoid fluorescence quenching of dye.

### Nanoparticle size determination

Nanoparticles size measurement was performed using a Zetasizer Nano ZS90 (Malvern Instrument, Westborough, MA.). Nanoparticles were resuspended into water and transferred into a disposable cuvette. Sample containing cuvette was placed into equipment and three measurements were taken to record the size of nanoparticles at 25 °C.

### Nanoparticles Zeta Potential

The surface charge on the nanoparticles was measured by Zetasizer ZS90 (Malvern Instrument, Westborough, MA.). In brief, 50 μl of chitosan nanoparticles sample was diluted 20 times and transferred into a specialized cuvette for zeta potential measurement. Specialized cuvette containing samples was placed into the sample holder and three measurements were taken at 25 °C.

### Surface Morphology Study

The surface morphology of CPX loaded nanoparticles was characterized by using a ZESS Scanning Electron Microscope (Tescan USA Inc. USA). Lyophilized nanoparticles with trehalose powders were mounted onto aluminum stubs using double sided adhesive tape. Samples were then made electrically conductive by coating in a vacuum with a thin layer of gold using a Polaron SC500 Gold Sputter Coater (Quotum technologies, Newhaven, UK). The coated specimen was then examined under the microscope operated at an acceleration voltage of 10 kV.

### Encapsulation Efficiency

The encapsulation efficiency of CPX loaded nanoparticles was determined using indirect method [13]. The encapsulation efficiency and loading capacity of nanoparticles were determined after collection of the nanoparticles by centrifugation at 12,000 rpm for 70 min at 4°C. The amount of free CPX in the supernatant was measured spectrophotometrically (Thermo Scientific) at 276 nm. The encapsulation efficiency (EE%) and the loading capacity (LC%) of the nanoparticles were calculated as follows:

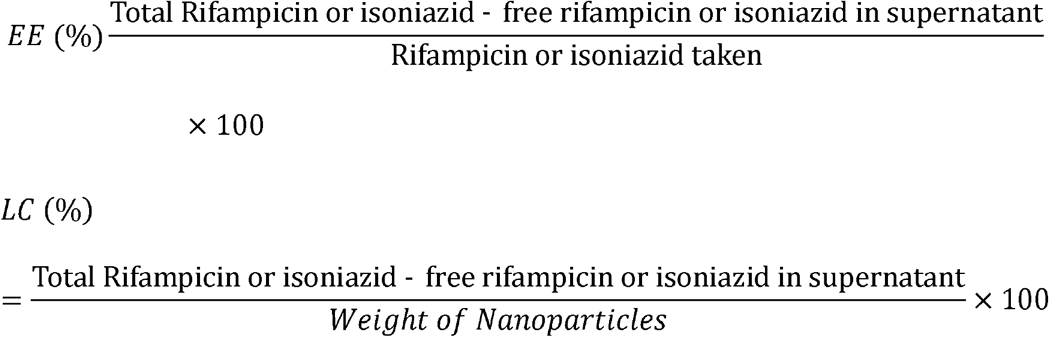

### Release Study

Ciprofloxacin release from nanoparticles was studied using a dialysis method. In brief, The CPX loaded chitosan nanoparticles were collected after centrifugation at 12,000 rpm for 70 min. The nanoparticles were transferred into a dialysis bag (containing the 5 mg equivalent of free CPX) with a cut-off molecular weight of 12 kDa and dialysis bag was immersed into 100 ml phosphate buffered saline at pH 7.4 or pH 5.2 to simulate body fluid pH & inside macrophage pH respectively at 37°C under continuous shaking. At predetermined time, 4 ml dialytic medium was taken out for analysis and an equal volume of freshly prepared PBS was added. The absorbance of the collected sample was taken at 276 nm by an UV–vis spectrometer. The released amount of CPX was quantified by referring to a calibration curve recorded from known amounts of CPX at the same condition.

### Uptake Study

Differentiated THP-1 cell line was used in uptake study. In brief, THP-1 cells were seeded in 6 well plate (0.1 x 106 cells/well) with 10 % FBS RPMI media, 20nM PMA and incubated at 37 °C for 36 hrs in a CO2 incubator (5% CO2 and 95% RH). After 36 hrs, the supernatant was removed, and fresh media was added to each well. The differentiated THP-1 cells were then incubated with FITC alone and FITC dye loaded nanoparticles for 6 hrs. Each well was washed three times with PBS. Cells were scraped using a scraper and cell suspension was analyzed for nanoparticles uptake study using Flow Cytometer (BD FACS Calibur, USA).

### Toxicity Assay

Nanoparticles induced cytotoxicity study was performed using different assay such as trypan blue dye exclusion assay, neutral dye uptake & accumulation assay and haemoglobin release assay on the different cells.

### Trypan blue dye exclusion test

The trypan blue dye exclusion test was performed according to the method of Tennant [14]. Failure to exclude trypan blue reflects a loss of plasma membrane integrity associated with necrosis [15]. Three replicates per concentration were maintained. Cells were mixed with equal volume of trypan blue stain and observed under a microscope and counted for stained cells.

### Neutral red uptake assay

The neutral red (NR) uptake assay was done to determine the accumulation of neutral red dye in the lysosomes of viable cells [16].The uptake of NR into the lysosomes/endosomes and vacuoles of living cells is used as a quantitative indication of cell number and viability. The neutral red retention assay was performed according to the methods described by Guillermo et al [17]. In brief, 3T3 fibroblasts cell line were seeded into 96-well plates (10,000 cells per well), and incubated overnight for cell recovery and adherence and for exponential growth. Cells were then treated with different concentrations (1, 10, 100 or 10000 μg/ml) of CS-NPs, CPX-CS NPs or with vehicle (PBS-buffer, pH 7.4) alone as a negative control for 24 hrs at 37°C under a humidified atmosphere of 5% CO2. After the incubation period the cells were washed carefully with PBS and incubated with the neutral red solution (4 μg/ ml in PBS) for 3 h. After incubation, cells were washed with PBS three times to remove all dye not incorporated within cellular lysosomes. A total of 150 μl of acidified ethanol (1% acetic acid, 50% ethanol, 49% distilled water) was added to each well and incubated in the dark for 15 min, before being read at 540 nm in the plate reader. All experiments were performed in triplicate. The cell viability (%) relative to the control wells containing cell culture medium without nanoparticles or PBS as a vehicle was calculated by the following equation:

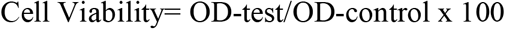

Where OD-test is the light absorbance (540 nm) of the test sample and OD-control is the light absorbance (540 nm) of the control sample. Each sample was measured in triplicate from three independent experiments.

### Hemoglobin Release Assay

Blood compatibility studies was performed with RBCs isolated from goat blood sample. In brief 10 ml of blood sample of the goat was obtained from a slaughter house in tube having anticoagulant EDTA after proper mixing and centrifuged at 1500 x g for 10 minutes. Red blood cells (RBCs) were collected and washed thrice with phosphate buffer solution (PBS) and spin for 7 min. at 1000g. RBC’s were resuspended into PBS and diluted up to 20% erythrocyte stock solution. The nanoparticles suspension was added to erythrocyte stock solution at different concentrations (1, 10, 100 or 1000 μg/ml) of CS-NPs, CPX-CS NPs or with vehicle (Normal saline) alone as a negative control for 2 hrs. The mixtures were incubated at 37□C in continuous shaking water bath for 2 hour. After the centrifugation at 1000 x g for 5 minutes, the supernatant absorbance recorded at 560 nm. The saline solution alone was used as negative control (0 % lysis) and 0.1 % triton in PBS used as positive control (100% lysis) [18]. The percent of hemolysis was calculated using the formula;

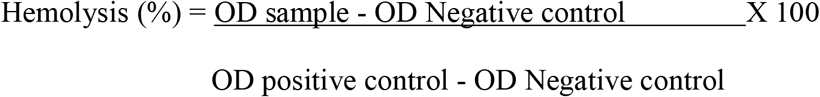

### Lipid Peroxidation Assay

Membrane damage in RBCs was determined by quantifying the release of malondialdehyde (MDA) after nanoparticles exposure. MDA is a product of lipid peroxidation and its release levels are indicative of membrane damage. Lipid peroxidation in erythrocytes was assayed as a method described by Stern et. al. [19]. In brief, RBCs incubated with CS NP, CPX-CS NP or CPX in equivalent amounts of drug in solution with 1ml of RBC (20%), 1ml of 10% ice cold trichloroacetic acid was added and vortexed. The mixture was centrifuged at 3000 rpm for 10 min followed by addition of 1 ml 0.67% w/v 2-thiobarbituric acid (in 0.1N NaOH) to 1ml supernatant which was kept in a boiling water bath for 10min. The final reaction mixture was cooled, diluted with 1ml distilled water and absorbance was recorded at 535nm. The results were expressed as nM MDA formed/ml erythrocytes. Molar extinction coefficient of MDA-TBA complex at 535 nm is 1.56X108/M/cm.

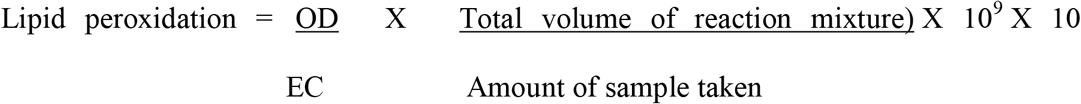

### Antibacterial activity

The antibacterial activity of CPX loaded chitosan nanoparticles was studied by agar diffusion assay method. In brief CPX-CS NPs containing 0.15 μg/ml equivalent of free CPX were taken in this study. A molten Mueller-Hinton agar stabilized at 45°C was seeded with 0.1 ml of a 24 h broth culture of the test organism (E. coli) containing approximately 105 cfu / ml in a sterile petri dish and allowed to set. Wells of 6mm diameter were created with a sterile cork borer and filled with 50 μl nanoparticles or free drug. The plates were pre-incubated for 1 h at room temperature to allow for diffusion of the solution and then incubated for 24 h. The zones of inhibition were measured (mean, n=3).

## Results

### Ciprofloxacin loaded nanoparticles characteristics

Ciprofloxacin loaded chitosan nanoparticles were prepared by gelation method by adding TTP. The CS/TPP weight ratio influenced the particle size and zeta potential of nanoparticles, by increasing in the CS/TPP weight ratio nanoparticle size and zeta potential was increased (195.6 nm to 229.1 nm and +24.86 to +28.35mV). The colloidal stability of formulations prepared was observed after 1 month storage in the refrigerator. Particle aggregation was measured in terms of increase in polydispersity and particles size of same particles at the different time was observed in one formulation of CS/TPP weight ratio 9/1 and not in 5/1 ratio, therefore formulation with 5/1 weight ratio was selected for further studies (300.8 nm and PDI was 0.350). Increase in chitosan concentration from 1 to 2% W/V affected the size and zeta potential of the optimized nanoparticles as shown in Table 1. The increase of CPX concentration led to a decrease of encapsulation efficiency. Nevertheless, CPX was entrapped in the matrix of nanoparticles to the appreciable extent (43–47%).

**Table 1.** Encapsulation Efficiency, Particle Size, Polydispersity Index and Zeta Potential of CPX-Loaded Chitosan Nanoparticles. (n=5)

| CS/TPP weight ratio | Ciprofloxacin Hydrochloride (mg/ml) | Average particle size (nm) | Polydispersity (PDI) | Zeta potential (mV) | Encapsulation efficiency (%) |
| --- | --- | --- | --- | --- | --- |
| 5:1(CS 1mg/ml) & TPP (0.5mg/ml)<br>**Formulation 1** | 0.5 | 195.6±11 | 0.249 | +24.86±1.2 | 43±2.29% |
|  | 1 | 185.5±2 | 0.217 | +24.91±0.6 | 42±2.35% |
| 5:1 (CS 2mg/ml & PP(1mg/ml)<br>**Formulation 2** | 0.5 | 214.8 ±8 | 0.249 | +29.6±0.8 | 46±2.15% |
|  | 1 | 252.9±9 | 0.254 | +27.0±0.7 | 43±2.27% |
| 9:1 (CS 1mg/ml & TPP 1mg/ml)<br>**Formulation 3** | 0.5 | 229.1±13 | 0.250 | +28.35±2.6 | 43±2.25% |
|  | 1 | 235.4±16 | 0.289 | +29.31±2.1 | 43±4.54% |

The surface morphology of CPX loaded NP was studied after lyophilization and trehalose used as cryoprotectant. In surface morphology study reveals size of particles around 250 nm and these particles are irregular in shape and sticky in nature. Particles are deposited on the trehalose iceberg (Fig. 1).

**Fig. 1.**
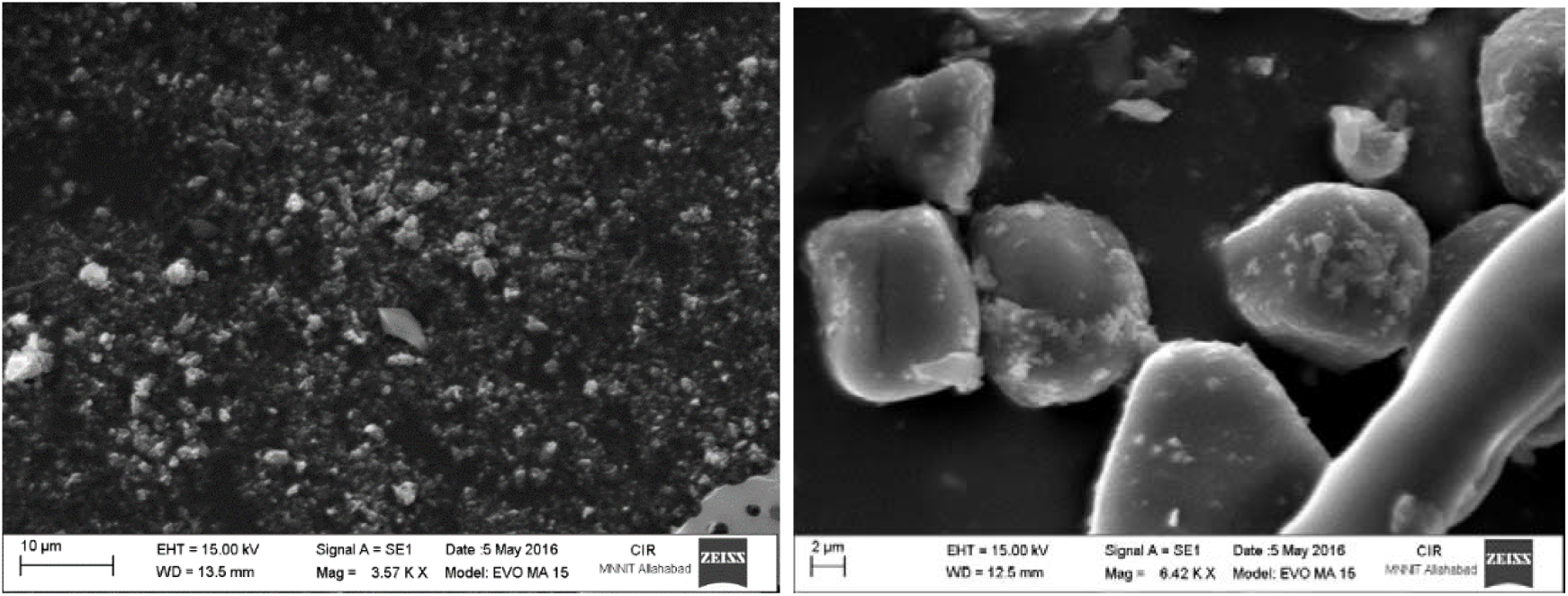
Surface Morphology of CPX loaded chitosan nanoparticles after lyophilization along with trehalose cryoprotectant using scanning electron microscopy (SEM) image analysis.

### Drug release and stability of chitosan nanoparticles

The release profile of CPX loaded chitosan nanoparticles are shown in Fig. 2, Where Ciprofloxacin in solution was released rapidly from the dialysis bag within the first 12 hrs. and from CPX loaded chitosan nanoparticles showed a biphasic pattern with an initial burst drug release followed by a more sustained release with about 1% of the drug is immediately released in 1 hour and 27% in 24 hrs. in pH 7.4 release medium. Similarly, at pH 5.2, CPX loaded nanoparticles, about 1% of the drug was immediately released in 1 hour and 13% in 24 hr.

**Fig. 2.**
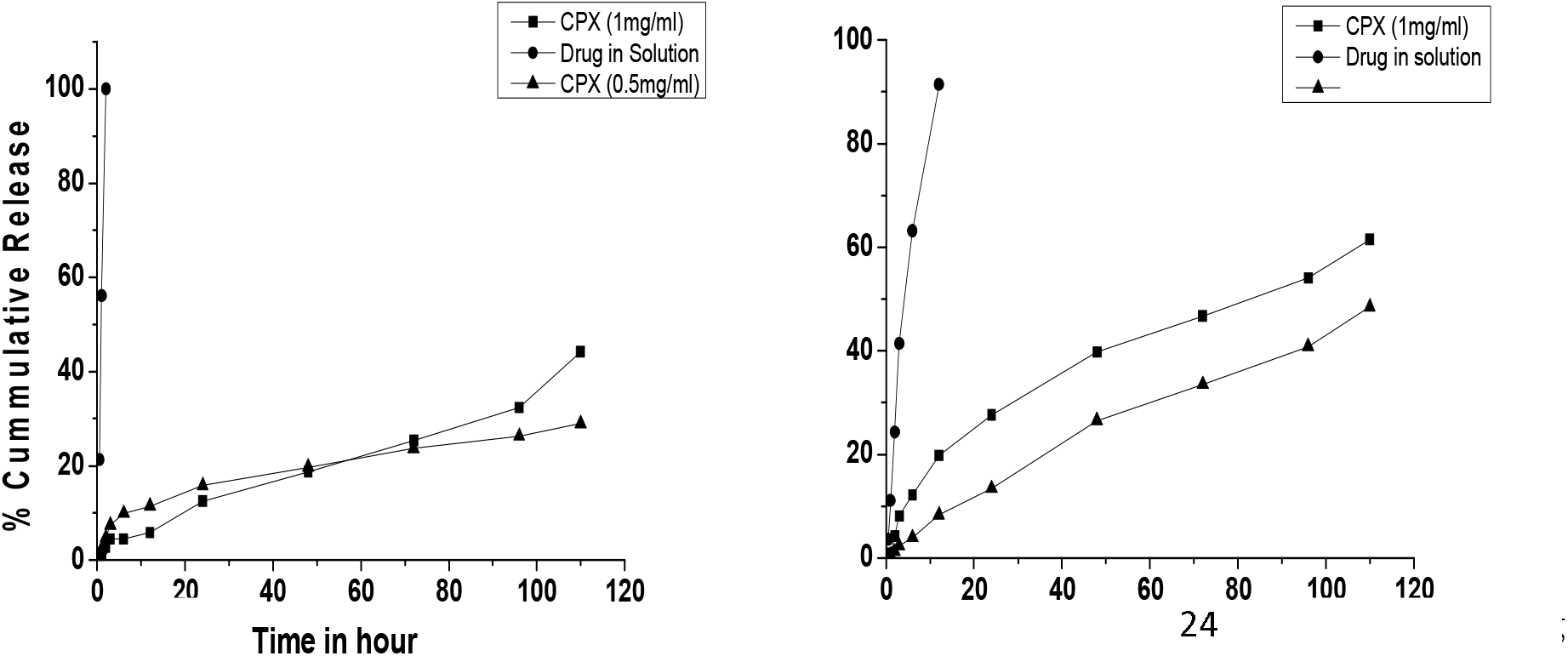
*In vitro* drug release study of Ciprofloxacin hydrochloride loaded Chitosan nanoparticles in **(A)** PBS buffer at pH 5.2 and **(B)** PBS buffer at pH 7.4

Stability of nanoparticles prepared by gelation method is an essential requirement for pharmaceutical applications, the storage stability and biocompatibility of the nanoparticles is a great concern. It is known that tiny particles are inclined to agglomerate with each other to reduce the surface area, and hence to reduce the free surface energy. Stability of prepared nanoparticles was analyzed at 4 °C and room temperature for 30 days. All the particles with 5/1 weight ratio were rather stable with negligible size fluctuation 201.6 nm and PDI-0.269, while particles with 9/1 weight ratio showed size increment and decrease in drug content (Table 2).

**Table 2.** Particle Size, Polydispersity Index and drug content of CPX-Loaded Chitosan Nanoparticles stored at room temperature after 30 days.

| CS/TPP weight ratio | Average particle size (nm) | Polydispersity (PDI) | Drug Content |
| --- | --- | --- | --- |
| 5:1 (CS 1mg/ml & TPP)<br>(0.5mg/ml) | 203.6±22 | 0.269 | 96% |
| 5:1 (CS 2mg/ml & TPP)<br>(1mg/ml) | 262.9±35 | 0.284 | 96.2% |
| 9:1 (CS 1mg/ml & TPP<br>1mg/ml) | 300.8±31 | 0.350 | 85% |

### Cellular uptake of FITC loaded chitosan nanoparticles

Bacteria reside into cells of the mammary gland, it’s very crucial for nanoparticles to inter inside cells for efficient killing of bugs inside affected cells. Uptake study of chitosan nanoparticles was studied in differentiated THP-1 cells by flow cytometry study (Figure 3). The uptake of FITC loaded nanoparticles by differentiated THP-1 cells was measured by flow cytometry, 10000 cells event for each sample was acquired. Positive and negative control was used as unlabeled and labelled cells to set the voltage of the detector for this experiment. In this study it was clearly shown FITC loaded NPs maximally taken by differentiated THP-1 cells.

**Fig. 3.**
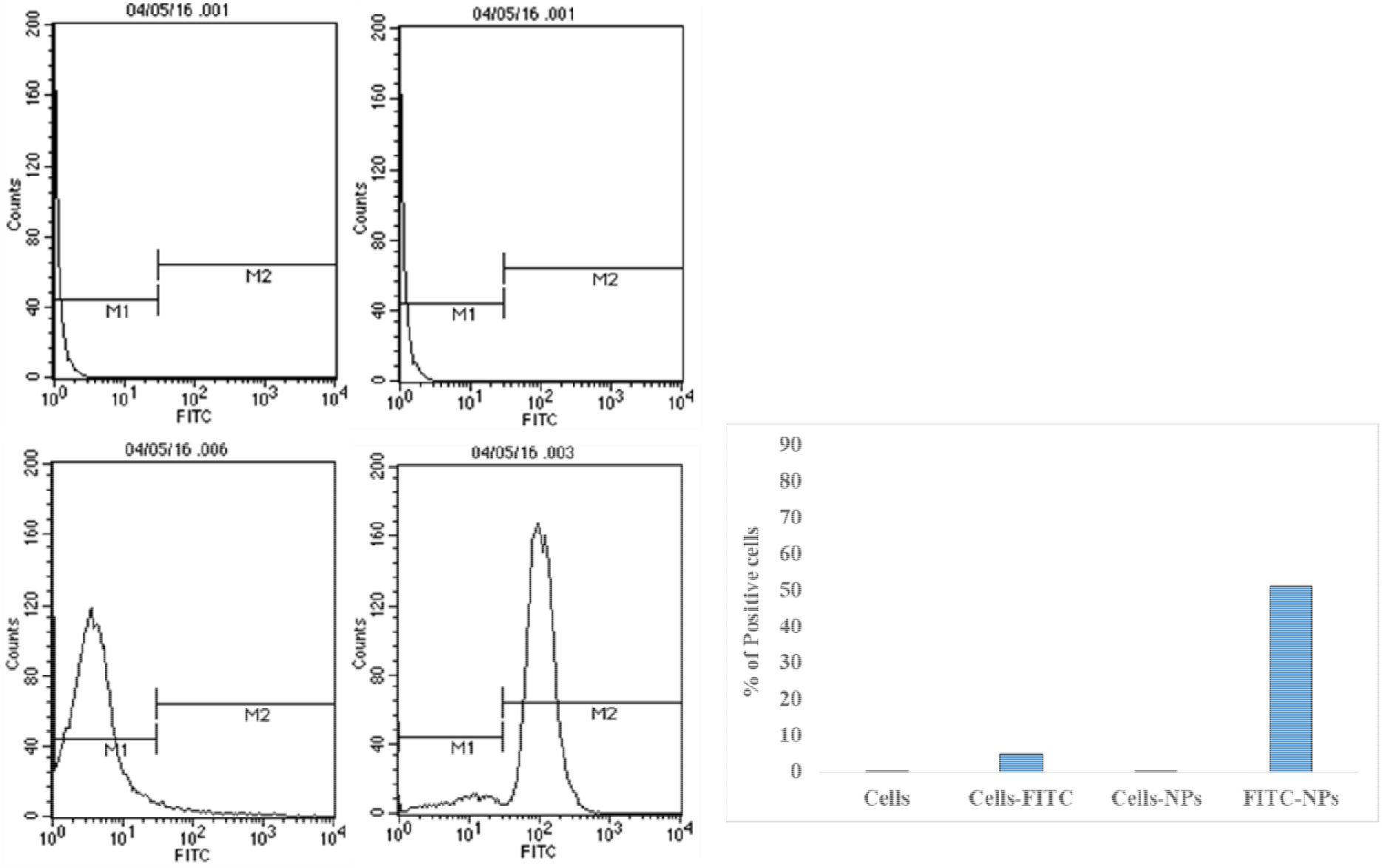
Cellular uptake of FITC dye loaded chitosan nanoparticles **A**. Histogram of cell and **B**. Percentage of cells uptake dye loaded nanoparticles by flow cytometry analysis. (n=3, ±SD)

### Antimicrobial activity of nanoparticles against clinical isolates

Antimicrobial activity of CPX loaded nanoparticles was evaluated against clinical isolates of E.coli. and S. aureus. by agar diffusion study (Figure 4). Efficacy of antimicrobial activity of CPX loaded nanoparticles is dose dependent, at higher amounts of nanoparticles show a high zone of inhibition on agar plate. It also observed antimicrobial activity against S. aureus is more efficient in comparison to E. coli.

**Fig. 4.**
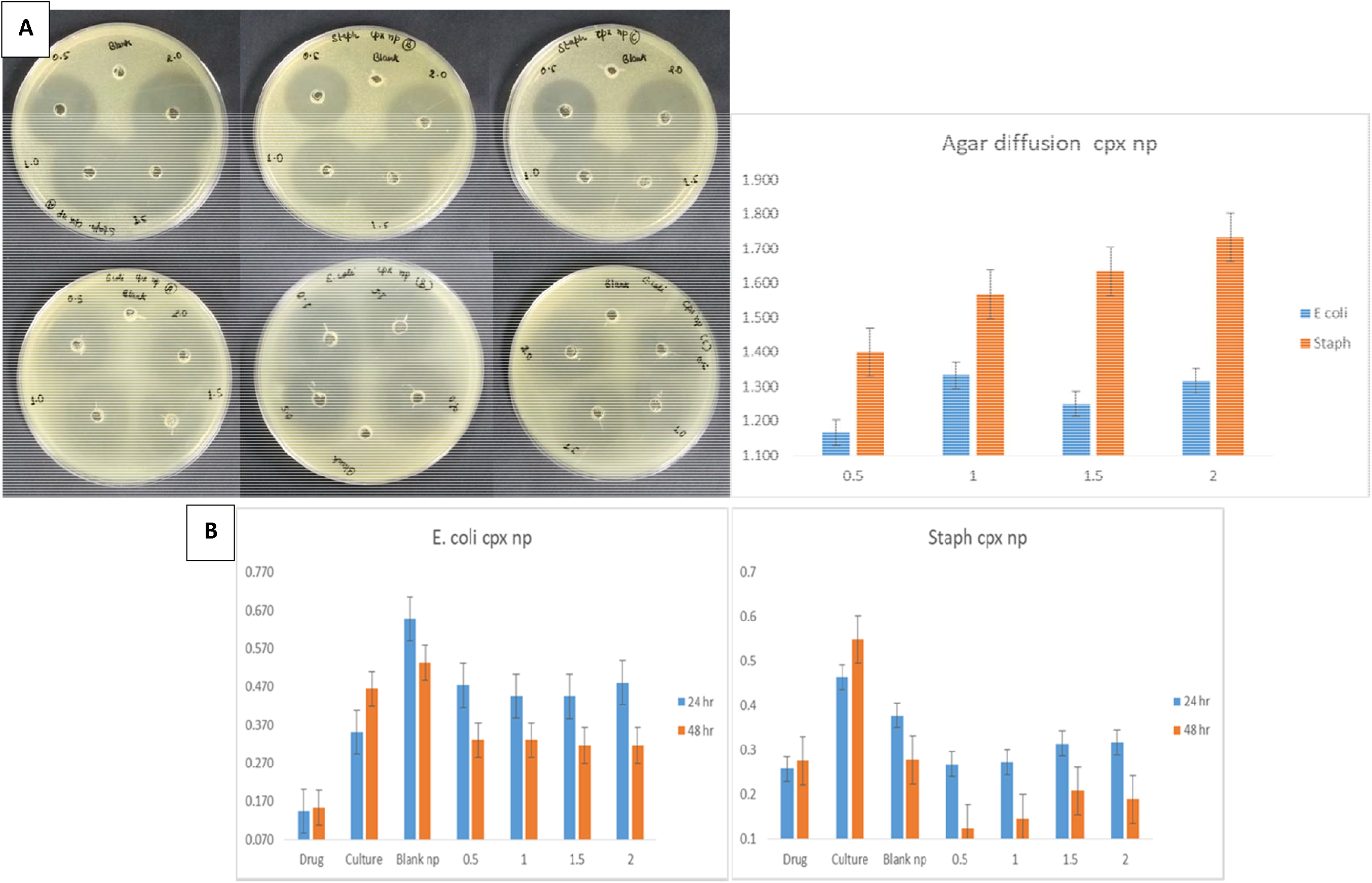
Antimicrobial activity of CPX loaded nanoparticles against clinical isolate *S. aureus* and *E. coli*. by **A**. agar diffusion assay and **B**. broth culture method at different dose of nanoparticles. (n=3, ±SD).

### Minimum inhibition concentration (MIC)

Growth inhibition of E.coli was studied in broth culture containing CPX or CPX-CS NPs respectively. No anti-microbial activity was observed with bare Chitosan NPs even at concentration ranges between 0 and 100 μg/ml. The MIC value of CPX alone was 0.16 μg/ml, and that of CPX-CS NPs was 0.12–0.16 μg/ml. This result indicates that the antimicrobial activity of CPX-CS NPs was not compromised during CPX loaded nanoparticles manufacturing method. MIC of CPX loaded nanoparticles comparable with CPX in solution and was not affected during the process of encapsulation into Chitosan nanoparticles.

### Cytotoxicity of CPX loaded Nanoparticles

In the preclinical development of a new pharmaceutical formulation, the cytotoxicity characterization is a one of critical parameters. We studied different toxicity induced parameters after exposure to the cells with nanoparticles such as Trypan blue dye exclusion assay, Neutral red uptake assay and Hemoglobin Release Assay for plasma membrane integrity of cells, viability of cells and blood cells compatibility assay with formulation.

In trypan blue dye exclusion assay we study the effect of CPX loaded NPs on membrane integrity of 3T3 fibroblast cell line. Relative to negative control, no statistically significant change was observed in the cell viability of cells in different treated groups with blank NPs, CPX loaded NPs (Fig. 5 A). In neutral red uptake study changes produced by toxic substances result in decreased uptake and binding of neutral red, making it possible to distinguish between viable, damaged or dead cells via spectrophotometric measurements. Alterations of the cell surface or the sensitive lysosomal membrane lead to lysosomal fragility and other changes that gradually become irreversible. Cells treated with blank NPs or CPX loaded NPs did not show any significant change of dye uptake with reference to control. We examined the viability of 3T3 fibroblast cultures treated with the different amounts of CPX in solution, CS-NP and CPX-CS NPs using the neutral red assay. The results showed that CPX in solution, CS-NP and CPX-CS NPs did not affect the viability of these cells at the doses used in the study. The cell viability plot (Fig. 5 C) showed that more than 91% cells were viable after 24 hr of incubation even in the presence of very high concentrations of CS-NP and CPX-CS nanoparticles (1000 μg/ml). No alteration of cell morphology of any of the cell lines was observed under the microscope. In addition, the cells were challenged up to 1 mg/ml of CS-NP and CPX-CS NPs and do not reduce the viability of cells, thus confirmed that no toxicity or minimal toxicity induced by CPX loaded nanoparticles are very low and the IC_50_ > 1 mg/ml. The results of the neutral red uptake assay did not indicate lysosomal fragility. The neutral red assay confirmed that CS-NP and CPX-CS NPs are non-toxic and are cytocompatible (figure 2). Hemocompatibility is an important parameter to access the biological safety of nanomaterials. As a preliminary biocompatibility study of formulation and their suitability as a formulation, we measured blood cell RBCs compatibility by exposing CPX loaded nanoparticles with RBCs (Fig. 5 B). No significant RBCs lysis was observed after exposure of different concentrations of CS-NPs, CPX-CS NPs, CPX in solution (less than 5% and comparable with the control). Lipid peroxidation for membrane damage assay

**Fig. 5.**
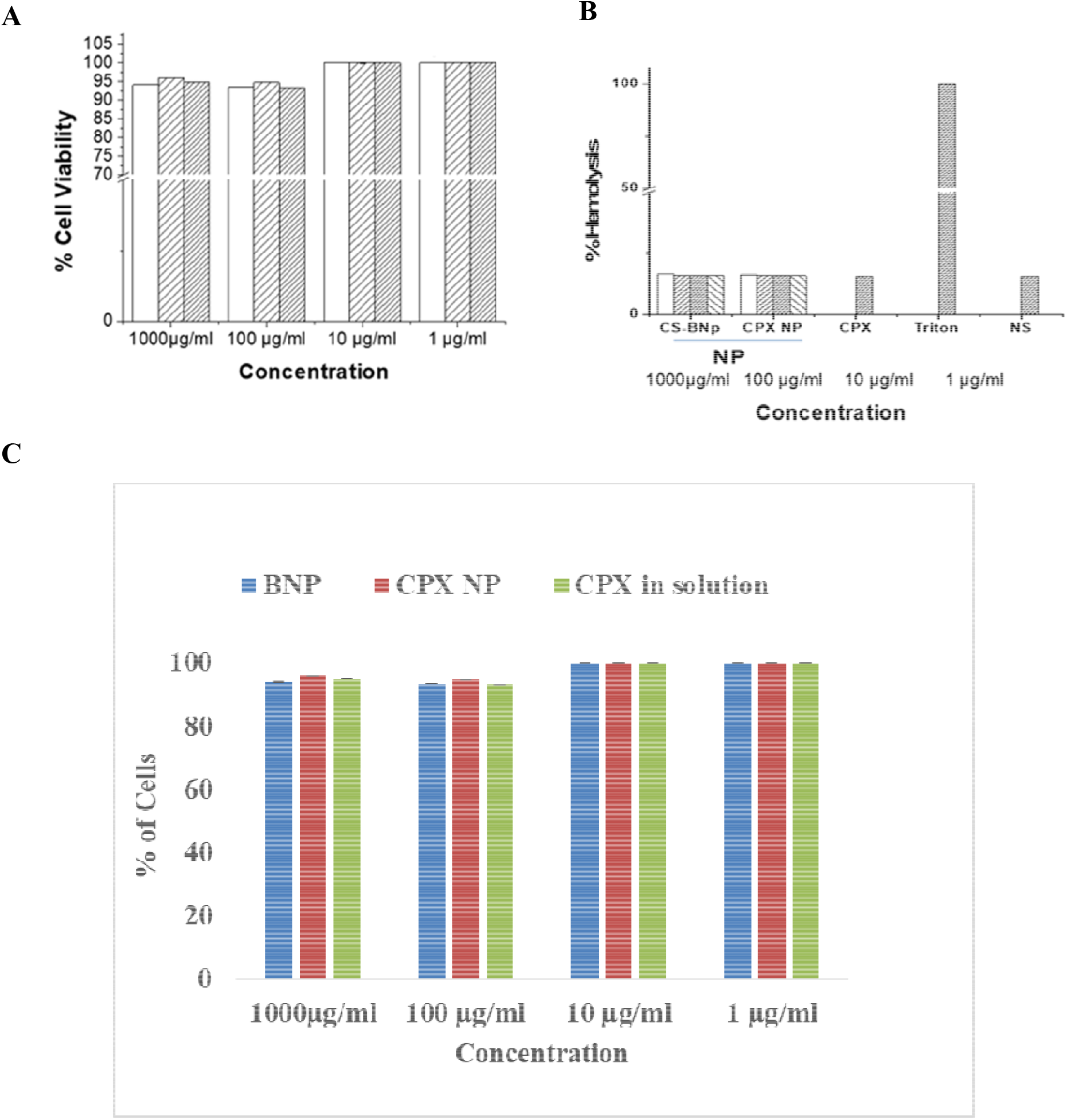
Toxicity assay after cells exposed with CPX loaded nanoparticles to the cells **(A)** The cell viability assay by trypan blue dye exclusion assay in 3T3 cells exposed with different dose of CPX, CPX-NPs or blank NPs (1, 10, 100, 1000 ug/ml) **(B)** Hemolysis assay to study membrane integrity assay to access membrane damage by CPX loaded NPs. **(C)** Neutral red dye uptake assay, cells exposed with ciprofloxacin (CPX), chitosan blank nanoparticles (BNP) and CPX loaded chitosan nanoparticles (CPX-CS NP) at different concentrations. 0.1% Triton X-100 as a positive control and normal saline (NS) used as negative control.

Oxidative degradation of cell membranes of cells after exposure with CPX loaded nanoparticles was measured. Membrane degradation is initiated by the presence of ROS. Oxidative degradation was measured after exposure of cell with CPX loaded nanoparticles by the presence of malondialdehyde or other thiobarbituric acid reactive substances (In Figure 6) [20]. It was shown that blank nanoparticles and CPX loaded nanoparticles induced a dose dependent peroxidative reaction of the PUFAs. In normal cells comparison with the control group, lipid peroxidation was increased, and drugs containing nanoparticles were capable of generating higher lipid peroxidation at 100 μg/ml and above concentration though the differences were not significant (p≥0.05). Higher ROS generation plays an important role in drug delivery as it plays an important role in host defence against microbes and also suggests activation of innate immunity. The lipid peroxidation is higher than the drug in solution which indirectly indicates beneficial effect of drug delivery and efficient intracellular killing bacteria in the infected host.

**Fig. 6.**
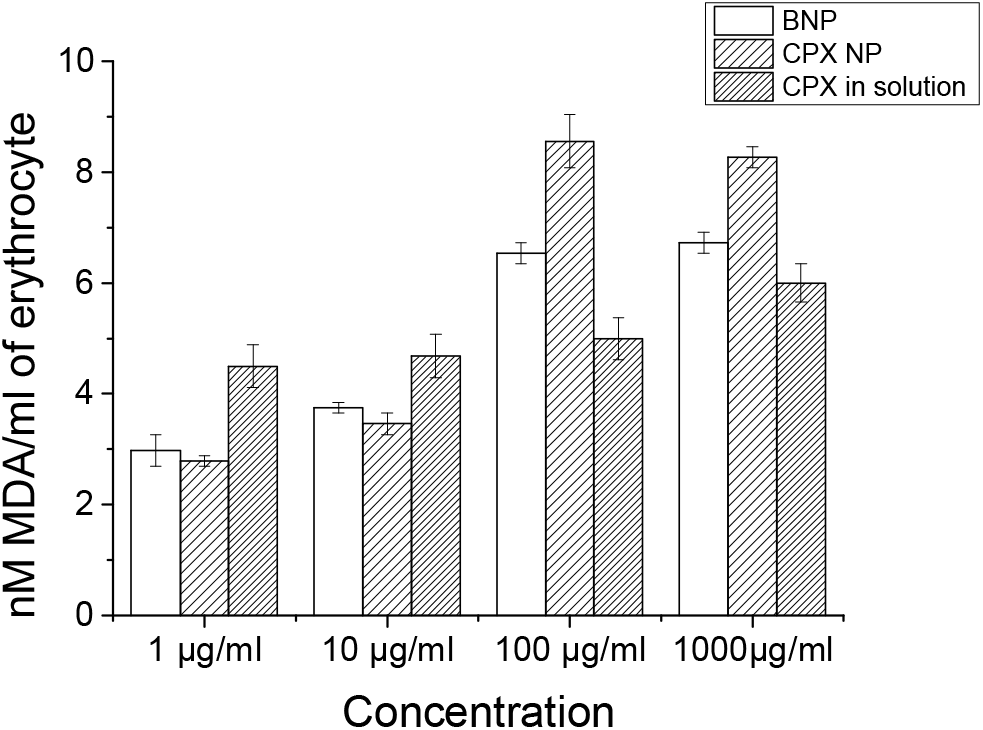
CPX loaded Chitosan nanoparticles exposure to the cells and membrane damage of cells access by lipid peroxidation assay (n=3, ±SD).

## Discussion

In milking dairy animal mastitis mainly caused by *E.coli.* and *S. aureus,* which contaminate milk as well as reduced milk production [21,22]. It’s very tough to treat mastitis disease by antibiotics treatments in milking animals due to milking at regular intervals, which interfere to maintain required concentration of antibiotics at the site of infection. Antibiotics loaded nanocarriers offer great advantages over conventional therapy such as targeted delivery on disease site and controlled release of drug which help to maintain optimal concentration of drug at affected site [23]. The present studies show CPX encapsulated into a chitosan based nanocarrier able to release drug at physiological and acidic pH in controlled manner, which can overcome limitations of antibiotics treatments of mastitis in dairy animals[24].

To overcome challenges posed to maintain sufficient concentration of antibiotics at disease site, we hypothesize controlled release of antibiotics from nanocarriers can address this problem. We encapsulated broad spectrum antibiotics CPX into nano-size chitosan particles. Antibiotics encapsulated nanosize particles release CPX at acidic and basic pH medium in controlled manner, up to 8 days it releases 60-70% of total drug[24]. Stability of formulation plays a very important role and also a very critical factor for the development of drug loaded nanocarriers for therapeutics use. We studied the stability of chitosan nanoparticles synthesized using different ratios of chitosan and TPP and found that 5:1 ratio stable upto 30 days in liquid formulation. We also found that these CPX loaded nanoparticles drug release is pH dependent. It can release drug faster at a higher pH than around lower pH (pH 7.4 > pH 5.2) [25]. The faster drug elution at neutral pH might be due to the swelling of the polymer matrix, which is brought about by deprotonation of amine group of chitosan. The drug release was increased with increase in chitosan concentration. The diffusion of drug from the surface creates a pore in the matrix which causes a channeling effect. Incorporation of higher concentration of drug causes more pore formation leading to faster and higher drug release from CPX loaded nanoparticles with higher encapsulation efficiency. In uptake study of these particles show easily uptaken by cells in in-vitro condition. We also performed efficacy study against clinical isolates of bacterium and CPX loaded particles show good efficacy in agar diffusion as well as in broth culture experiments. In cytotoxicity study we used RBCs to incubate with CPX loaded nanoparticles and found to be no or with minimal toxicity in comparison to CPX drug alone. We performed different toxicity assay such as MTT assay, Hemolysis assay & lipid peroxidase assay.

## Conclusions

Our data shows that CPX loaded chitosan nanoparticles can provide a controlled drug release which depend on the pH of surrounding tissue. CPX loaded into nanoparticles can kill bacteria efficiently. CPX loaded chitosan nanoparticles can be used for effective treatments of mastitis in dairy animals.

## Acknowledgments

We are thankful to the Department of Biotechnology (DBT) New Delhi, India for providing financial support, Research Fellowships to PY & ZIH from DBT. This work was supported by the Grant No. BT/PR21547/NNT/ 28/ 1232/2017 of Department of Biotechnology (DBT), Government of India.

## References

1. Fetrow J, Mann D, Butcher K, McDaniel B (1991) Production losses from mastitis: carry-over from the previous lactation. Journal of dairy science 74 (3):833–839. doi:10.3168/jds.S0022-0302(91)78232-5

2. Houben EH, Dijkhuizen AA, Van Arendonk JA, Huirne RB (1993) Short- and long-term production losses and repeatability of clinical mastitis in dairy cattle. Journal of dairy science 76 (9):2561–2578. doi:10.3168/jds.S0022-0302(93)77591-8

3. Mungube EO, Tenhagen BA, Regassa F, Kyule MN, Shiferaw Y, Kassa T, Baumann MP (2005) Reduced milk production in udder quarters with subclinical mastitis and associated economic losses in crossbred dairy cows in Ethiopia. Tropical animal health and production 37 (6):503–512

4. Dogan B, Klaessig S, Rishniw M, Almeida RA, Oliver SP, Simpson K, Schukken YH (2006) Adherent and invasive Escherichia coli are associated with persistent bovine mastitis. Veterinary microbiology 116 (4):270–282. doi:10.1016/j.vetmic.2006.04.023

5. Varella Coelho ML, Santos Nascimento JD, Fagundes PC, Madureira DJ, Oliveira SS, Vasconcelos de Paiva Brito MA, Freire Bastos Mdo C (2007) Activity of staphylococcal bacteriocins against Staphylococcus aureus and Streptococcus agalactiae involved in bovine mastitis. Research in microbiology 158 (7):625–630. doi:10.1016/j.resmic.2007.07.002

6. Chaneton L, Tirante L, Maito J, Chaves J, Bussmann LE (2008) Relationship between milk lactoferrin and etiological agent in the mastitic bovine mammary gland. Journal of dairy science 91 (5):1865–1873. doi:10.3168/jds.2007-0732

7. Petros RA, DeSimone JM (2010) Strategies in the design of nanoparticles for therapeutic applications. Nature reviews Drug discovery 9 (8):615–627. doi:10.1038/nrd2591

8. Zhang L, Pornpattananangku D, Hu CM, Huang CM (2010) Development of nanoparticles for antimicrobial drug delivery. Current medicinal chemistry 17 (6): 585–594

9. Gao P, Nie X, Zou M, Shi Y, Cheng G (2011) Recent advances in materials for extended-release antibiotic delivery system. The Journal of antibiotics 64 (9):625–634. doi:10.1038/ja.2011.58

10. Mascellino MT, Farinelli S, Iegri F, Iona E, De Simone C (1998) Antimicrobial activity of fluoroquinolones and other antibiotics on 1,116 clinical gram-positive and gram-negative isolates. Drugs under experimental and clinical research 24 (3):139–151

11. Grillon A, Schramm F, Kleinberg M, Jehl F (2016) Comparative Activity of Ciprofloxacin, Levofloxacin and Moxifloxacin against Klebsiella pneumoniae, Pseudomonas aeruginosa and Stenotrophomonas maltophilia Assessed by Minimum Inhibitory Concentrations and Time-Kill Studies. PloS one 11 (6):e0156690. doi:10.1371/journal.pone.0156690

12. Fan W, Yan W, Xu Z, Ni H (2012) Formation mechanism of monodisperse, low molecular weight chitosan nanoparticles by ionic gelation technique. Colloids and surfaces B, Biointerfaces 90:21–27. doi:10.1016/j.colsurfb.2011.09.042

13. Honary S, Ebrahimi P, Hadianamrei R (2014) Optimization of particle size and encapsulation efficiency of vancomycin nanoparticles by response surface methodology. Pharmaceutical development and technology 19 (8):987–998. doi:10.3109/10837450.2013.846375

14. Tennant JR (1964) Evaluation of the Trypan Blue Technique for Determination of Cell Viability. Transplantation 2:685–694

15. Bonfoco E, Krainc D, Ankarcrona M, Nicotera P, Lipton SA (1995) Apoptosis and necrosis: two distinct events induced, respectively, by mild and intense insults with N-methyl-D-aspartate or nitric oxide/superoxide in cortical cell cultures. Proceedings of the National Academy of Sciences of the United States of America 92 (16):7162–7166. doi:10.1073/pnas.92.16.7162

16. Borenfreund E, Puerner JA (1985) Toxicity determined in vitro by morphological alterations and neutral red absorption. Toxicology letters 24 (2-3): 119–124. doi:10.1016/0378-4274(85)90046-3

17. Repetto G, del Peso A, Zurita JL (2008) Neutral red uptake assay for the estimation of cell viability/cytotoxicity. Nature protocols 3 (7): 1125–1131. doi:10.1038/nprot.2008.75

18. Ravikumara NR, Madhusudhan B, Nagaraj TS, Hiremat SR, Raina G (2009) Preparation and evaluation of nimesulide-loaded ethylcellulose and methylcellulose nanoparticles and microparticles for oral delivery. Journal of biomaterials applications 24 (1):47–64. doi:10.1177/0885328209103406

19. Potter TM, Neun BW, Stern ST (2011) Assay to detect lipid peroxidation upon exposure to nanoparticles. Methods in molecular biology 697:181–189. doi:10.1007/978-1-60327-198-1_19

20. Yang XF, Yang YN (1997) Protective effects of calcium antagonists on cadmium-induced toxicity in rats. Biomedical and environmental sciences: BES 10 (4):402–407

21. Pang M, Xie X, Bao H, Sun L, He T, Zhao H, Zhou Y, Zhang L, Zhang H, Wei R, Xie K, Wang R (2018) Insights Into the Bovine Milk Microbiota in Dairy Farms With Different Incidence Rates of Subclinical Mastitis. Frontiers in microbiology 9:2379. doi:10.3389/fmicb.2018.02379

22. Fahim KM, Ismael E, Khalefa HS, Farag HS, Hamza DA (2019) Isolation and characterization of E. coli strains causing intramammary infections from dairy animals and wild birds. International journal of veterinary science and medicine 7 (1):61–70. doi:10.1080/23144599.2019.1691378

23. Algharib SA, Dawood A, Xie S (2020) Nanoparticles for treatment of bovine Staphylococcus aureus mastitis. Drug delivery 27 (1):292–308. doi:10.1080/10717544.2020.1724209

24. Barbosa AI, Costa Lima SA, Reis S (2019) Application of pH-Responsive Fucoidan/Chitosan Nanoparticles to Improve Oral Quercetin Delivery. Molecules 24 (2). doi:10.3390/molecules24020346

25. Wang Y, Qin F, Tan H, Zhang Y, Jiang M, Lu M, Yao X (2015) pH-responsive glycol chitosan-cross-linked carboxymethyl-beta-cyclodextrin nanoparticles for controlled release of anticancer drugs. International journal of nanomedicine 10:7359–7369. doi:10.2147/IJN.S91906

